# Visuospatial attention revamps cortical processing of sound: restrict stimulus uncertainty

**DOI:** 10.1101/2020.07.30.229948

**Authors:** F. Cervantes Constantino, T. Sánchez-Costa, G. A. Cipriani, A. Carboni

**Affiliations:** Centro de Investigación Básica en Psicología; Universidad de la República; Montevideo, Uruguay; Instituto de Fundamentos y Métodos en Psicología; Universidad de la República; Montevideo, Uruguay; Instituto de Investigaciones Biológicas “Clemente Estable”; Montevideo, Uruguay; Facultad de Psicología, Universidad de Granada; Granada, Spain

**Author notes:** **Corresponding author:** Francisco Cervantes Constantino, Ph.D., Centro de Investigación Básica en Psicología, Facultad de Psicología, Universidad de la República, Dr. Tristán Narvaja 1674, Montevideo 11200 Uruguay.

## Abstract

Selective attentional biases arising from one sensory modality may manifest in another. The effects of visuospatial attention, often considered a foundation for visual object perception, are unclear in the auditory domain during audiovisual (AV) scene processing. This study investigates temporal and spatial factors that facilitate such cross-modal bias transfer at the neural level. Auditory encoding of random tone pips in AV scenes was investigated via a temporal response function model (TRF) of the participants’ electroencephalogram (*N*=30). The spatially uninformative pips were associated with spatially distributed visual contrast reversals (‘flips’) through asynchronous, probabilistic AV temporal onset distributions. Participants deployed visuospatial selection on these AV stimuli to perform a task. A late (~300 ms) cross-modal transfer of the unimodal attentional bias was found on the neural representation of pips. Transfer depended on the selected visual input being (*i*) presented during or shortly after a related sound in a relatively limited temporal window (<165 ms); and (*ii*) positioned across limited (1:4) visual foreground to background ratios. In addition, the magnitude of attentional enhancement was proportional to the proximity of flips to the foreground area. The results indicate that ongoing neural representations of sounds can incorporate relevant visuospatial attributes for auditory stream segregation.

## 1. Introduction

To parse scenes adequately, one must shift focus from one place to another as the relative importance of locations may change over time. Prevalent visuospatial coding in the human brain frames domains across and beyond vision (Groen et al., 2021). In human attentional systems, it is unclear how visuospatial biases transfer efficiently into sensory modalities such as hearing. The capacity of transfer has been addressed in the last decade with a focus on stimulus features (Koelewijn et al., 2010; Santangelo & Macaluso, 2012), especially by consistent observations of audiovisual (AV) integration (Busse et al., 2008; Degerman et al., 2007; Spence & Squire, 2003; Holmes & Spence, 2005; Roseboom et al., 2009). In the most prominent examples, driven by AV temporal proximity, ambiguous yet identical speech sounds bias a listener according to matching visual face displays. The examples include the ventriloquist illusion, by apparent spatial location shifts, and in the McGurk effect by changes to inferred speech utterances (Andersen et al., 2009; McGurk & MacDonald, 1976; Miller & D’Esposito, 2005; van Atteveldt et al., 2007).

What stimulus features determine the ability for visuospatial biases to influence auditory scene processing? Clearly among them is *timing,* i.e. the temporal coherence across the individual dynamics of both visual and auditory unimodal streams (Bizley et al., 2016; Spence & Frings, 2020; Talsma et al., 2010). Segregation of two simultaneous auditory streams can be promoted when only one is coherent with the visual scene dynamics (Maddox et al., 2015). Early facilitatory mechanisms include cortico-cortical cross-sensory interactions, some of which are independent of attention (Atilgan et al., 2018; Talsma & Woldorff, 2005). A second candidate appears to be *spatial* alignment between the auditory and visual streams (Fleming et al., 2020). Such factor appears less optimal to favor integration, however (Noesselt et al., 2005). For example, a light flash intermittent sound beep alarm may require additional time to be recognized as one same source, if poorly coordinated. Importantly, as vision dominates human spatial perception while hearing may do so for time (Burr et al., 2009), it is necessary to address the role of unimodal visuospatial factors alone for optimal cross-modal transfer. In the alarm example, given that the signals are well-aligned spatially, the most efficient search strategy may be to initially constrain the visual space of plausible locations. Foreground size is a determinant feature of visuospatial selection, yet it is still unknown how it contributes to the cross-modal transfer of visual biases onto the neural encoding of sound.

In addition, naturalistic conditions promoting robust multimodal perception often involve integration windows rather that temporally aligned signals (van Wassenhove et al., 2007). Importantly, asynchronous input implies that events from one modality may lead or lag those in another. Since bimodal asynchrony entails temporal order differences across unimodal events, this leads to *precedence* as a third basic factor to consider in the effective transfer of unimodal visuospatial biases. The first alternative set by this factor is that visual input best influences processing of subsequent auditory input through visually cued auditory selection (visual priming). The other alternative is that any auditory input establishes a temporal window, and visuospatially biased signals emerging within are most optimally incorporated into the course of such auditory processes (visual updating).

We investigate the hypothesis that precedence, and temporal and spatial window factors interact in the cross-modal transfer of visuospatial biases that promote cortical segregation of auditory streams. Here, diverse synthetic visual ‘dartboard’ stimuli featured a sequence of spatially distributed, local contrast reversals (‘flips’) at random times. The accompanying audio streams were composed of pure tones asynchronously paired one-to-one with a flip in the sequence. In any stimulus, individual pips randomly led or lagged flips in equal proportions (*precedence*), and its probabilistic asynchrony distribution was controlled (*AV temporal window*). To perform the task, human participants followed events across the full dartboard, or alternatively across half or quarter sectors of it, thereby attending to foregrounds of controlled size (*visuospatial window*). Using electroencephalographic (EEG) recordings, subjects’ auditory processing was then addressed via the temporal response function (TRF) model (Crosse et al., 2016; David et al., 2007). The auditory TRF represents the linear mapping between the temporal dynamics of sound input and that of the unfolding neural response time series. It has been related to the receptive field in neural populations given its ability to map rapid cortical encoding dynamics (Gaucher et al., 2012; Gourévitch et al., 2009). In attentional manipulations, TRFs estimated from foreground- versus background-related auditory input are often contrasted to measure selective enhancement/attenuation effects. Attentional effects measured by means of this technique are often observed in the relative changes to TRF peaks associated to each of the competing auditory source (Brodbeck et al., 2020; Ding & Simon, 2012; O’Sullivan et al., 2015). Here, we hypothesize the emergence of attentional effects subject to auditory encoding peak modulations, and the intervention of spatiotemporal distributional properties of stimulus input. For this, first we localized the presence of an attentional effect, and subsequently examined its relation to the interaction between precedence and temporal/spatial window conditions, thus promoting the neural segregation of auditory input by spatially selected visual input.

## 2. Method

### 2.1 Subjects

Thirty subjects (21 female; mean age 23.9 ± 4.0 SD) with no history of neurological or psychiatric disorder voluntarily participated in the study. All provided formal written informed consent. They reported normal hearing and normal or corrected to normal visual acuity. All experiments were performed in accordance with the WMA Declaration of Helsinki (World Medical Association, 2009) guidelines. The School of Psychology Research Ethics Committee at Universidad de la República approved the experimental procedures.

### 2.2 Setup and stimuli

#### 2.2.1 Visual stream

The visual part of the AV stimulus consisted of a sequence of non-simultaneous visual transients (Van der Burg et al., 2010) presented across a black and white dartboard-like disc display with black background. The disc subtended a radius of 23 degrees of visual angle and was segmented into a checkerboard pattern of 20 equally spaced concentric and 48 angular divisions. As in Capilla et al. (2016), the divisions were grouped into sectors defined by 4-by-4 checkerboard element divisions. The resulting 60 sectors were distributed over 5 eccentricity levels and 12 angular sections (Figure 1A). After 1 s, a single sector may undergo one instantaneous polarity reversal across all its internal element divisions, here termed a ‘flip’. Flips remained static unless stimulated again later, and their locations were uniformly balanced across eccentricities and disc quadrants. The time between flips ranged from 100 to 900 ms (see *Supplementary Method* for the distribution), and the final configuration was kept fixed for 0.5 s. The number of flips in a given video sequence depended on attentional condition (see 2.2.4 *Attentional conditions*, below). Visual stimuli were constructed with the MATLAB® software package (Natick, United States), and stored in .avi format. Presentation and response time logging were performed with PsychoPy (Peirce, 2007). Visual displays were delivered over a CRT monitor (E. Systems, Inc., CA) with 40 cm size, 83 dpi resolution, and 60 Hz refresh rate.

**Figure 1.**
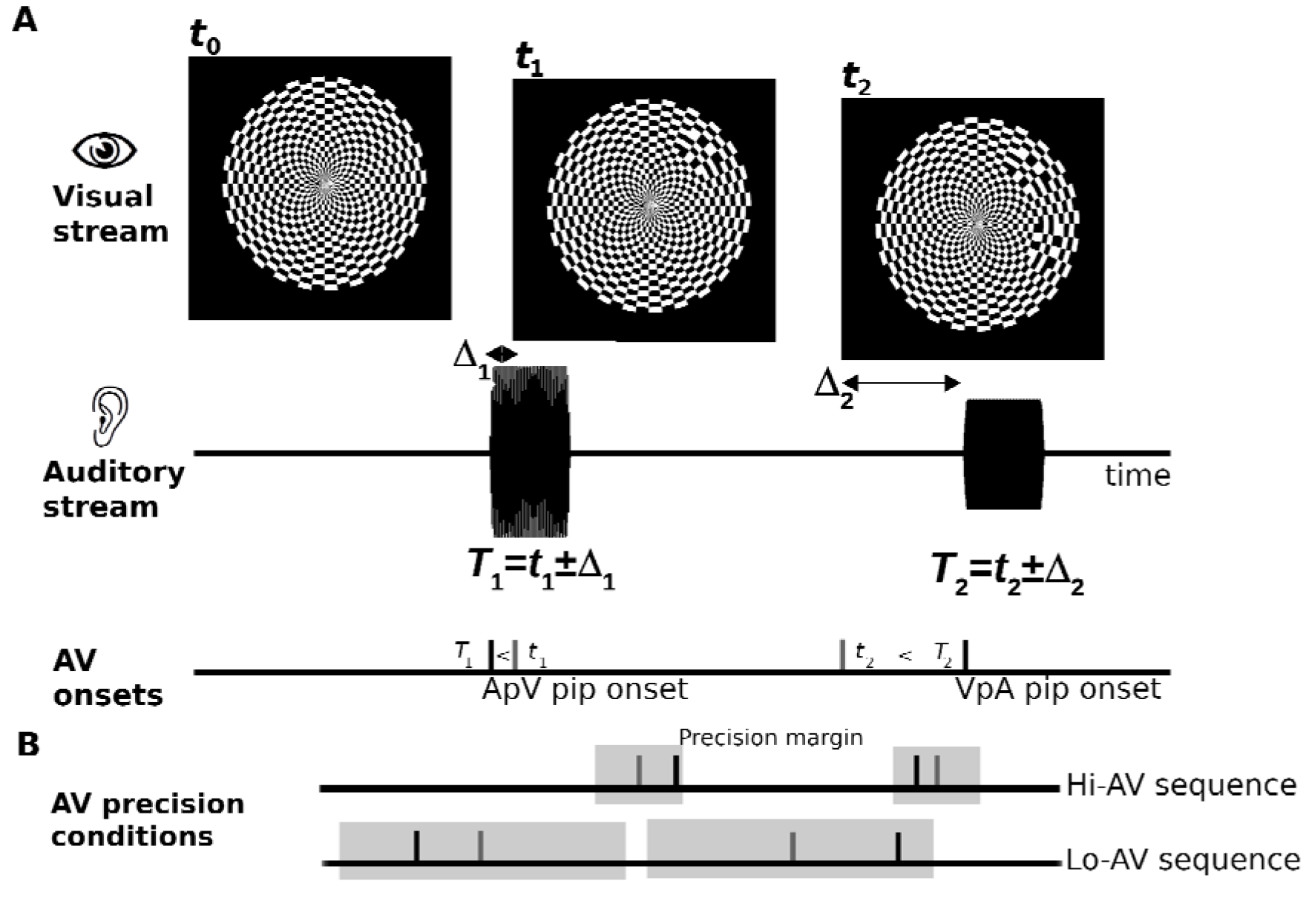
Basic audiovisual sequence and AV precision conditions. (A) Each audiovisual stimulus consisted of a dartboard disc displaying a series of local contrast changes or flips (‘V’), accompanied by tone pip sounds (‘A’). In this example, the first flip occurs in the upper right quadrant at time *t*_1_ and in the lower right at *t*_2_ (top). For each V, a corresponding A is presented with a positively or negatively delayed onset time *T_i_* (A precedes V ‘ApV’, middle; alternatively ‘VpA’, bottom). (B) The allowed range of AV asynchronies Δ_*i*_ defines the AV temporal window (or precision condition) for a given stimulus sequence. Grey areas illustrate different AV precision conditions at different AV stimulus sequences. High AV (‘Hi-AV’) sequences have narrower asynchrony margins (top) than do low AVsequences (‘Lo-AV’, bottom).

#### 2.2.2 Auditory stream

The accompanying audio presentation consisted of a sequence of pure tone pip sounds of 100 ms duration each (Figure 1A). Pip frequencies were selected from a fixed pool of 15 values ranging between 100 and 4846 Hz, separated by 2 equivalent rectangular bandwidth steps (Glasberg & Moore, 1990), and pip frequency values were equally balanced for each auditory sequence. Pips were modulated with 5 ms raised cosine on- and off-ramps. Pip amplitudes were calibrated according to the 60-phon normal equal-loudness-level contour (ISO 226:2003) to adjust for perceived relative loudness differences across frequencies. The number of pips in a given audio sequence equaled that of the flips in the visual stream. Each tone onset time was defined according to the nominal time of the corresponding flip in the visual stream plus a uniformly distributed random shift (Figure 1A, *Supplementary Method*). The distribution had zero mean and its bounds were determined by the AV precision condition for that stimulus sequence, as indicated in 2.2.3 *Audiovisual stimuli,* below. Unlike flips, pips were allowed to overlap in time. Individual pip frequencies were not associated with the angular position of corresponding flips (see *Supplementary Method* for eccentricity control). All auditory stimuli were constructed with MATLAB® at a sampling rate of 22.05 KHz in .wav format.

#### 2.2.3 Audiovisual stimuli

For each AV stimulus’ video sequence and corresponding audio stream, a fixed range of permitted asynchronies Δ_*i*_, between individual flips and pip onsets (Figure 1B) was used. Ten different ranges were defined as ±33 ms, ±66 ms, ±99 ms, and so forth up to ±330 ms temporal window asynchronies. Two hundred AV stimuli, equally distributed across the ten ranges (see below), were built by merging individual video and sound files into multimedia with Simulink® and transcoding into .m4v format with Video Converter (V3TApps / The HandBrake Team).

#### 2.2.4 Attentional conditions

AV stimuli were built for each attentional condition as follows: “Attend-All” (AA) stimuli consisted of a sequence of 15 pip-flip pairs, lasting about 4 – 5 s. These stimuli were intended for attending to the whole dartboard as foreground. “Attend-Half” (AH) stimuli consisted of a sequence of 30 pip-flip pairs, lasting about 7 – 8 s. These stimuli were intended for attending to either the upper or lower dartboard hemifield as foreground, ignoring visual events from the opposite hemifield. “Attend-Quarter” (AQ) stimuli consisted of a sequence of 60 pip-flip pairs, lasting about 12 – 15 s. These stimuli were intended for attending to either the upper-, lower-, -right or -left dartboard quadrant as foreground, ignoring visual events from all other three quadrants. These conditions ensured that the visuospatial windows determining foreground size consisted of 15 pip-flip pairs across all stimuli. The fixed pool of 200 AV stimuli was distributed as 40 AA, 80 AH and 80 AQ stimuli.

### 2.4 Task

The experiment was organized as a two-interval forced choice task based on a visuospatial attention manipulation. During initial fixation, participants were cued on which sector to attend to in the trial. Cues represented: AA by a full circle, or AH by an upwards/downwards triangle, or AQ by a diagonally tilted triangle pointing to a specific quadrant (Figure 2, inset). Two different AV stimuli from the cued attentional condition were subsequently presented in series (Figure 2, Video 1). The task for participants was to compare between the first and second stimulus foregrounds, for their overall sense of AV synchronicity between presentations. Participants were thereafter prompted: “Which sequence was more [less] coordinated?” (Figure 2). Subjects pressed a left/right arrow button to highlight their choice and then hit enter to validate. A feedback cue was then presented and the experiment ended after 100 trials. The full experimental sessions (*Supplementary Method*) lasted ~2 h. As compensation for their time, participants received a chocolate bar or a cinema ticket.

**Figure 2.**
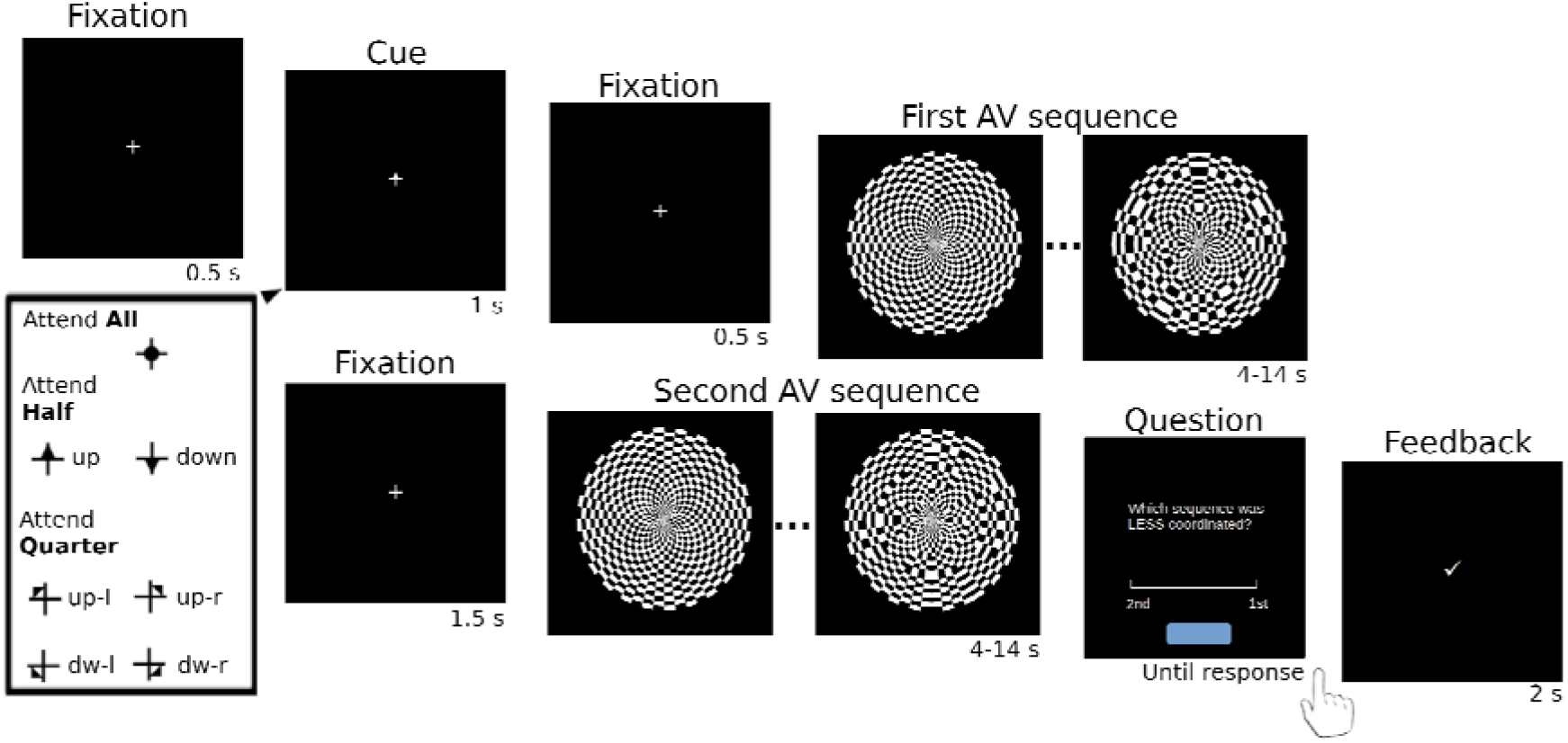
Two-interval forced choice task with AV stimuli. Participants were asked to compare two serially-presented AV sequences in an attentional task. An initial cue (inset) indicated the required foreground for the trial. Participants were then shown two AV sequences and deployed visuopatial attention over the same required foreground sector for both stimuli. In Attend-All, the foreground equaled the entire disc (full circle cue); for Attend-Half, only the upper/lower half (up/down triangle cues), and for Attend-Quarter, only one quadrant (tilted triangle cues). Participants had to compare the level of AV temporal precision of both stimuli foregrounds. Afterward, they were asked for a response and trial feedback was shown upon validation.

### 2.5 Data analysis

#### 2.5.1 EEG collection and pre-processing

Continuous EEG recordings were performed using a BioSemi ActiveTwo 64-channel system (BioSemi, The Netherlands) with 10/20 layout at 2048 Hz digitization rate with CMS/DRL (ground). Data analysis was performed in MATLAB 2018b offline. EEG subject data resampled at 1024 Hz were common average-referenced to the 64 scalp channels, after which DC offset was removed. Signals were bandpass-filtered between 1 and 40 Hz with a 20-order elliptical filter of 60 dB attenuation and 1 dB passband ripple. Trial recordings were epoched from -1 s to: 8 s relative to stimulus onset (AA), 11 s (AH), or 17 s (AQ). Epochs were then downsampled to 256 Hz and denoised (*Supplementary Method*). To improve SNR of data reflecting reproducible auditory activity specific to the processing of tones in the present stimuli, a spatial filter was constructed. The denoising by spatial filtering (DSS) (de Cheveigné & Parra, 2014; de Cheveigné & Simon, 2008b) data-driven procedure was applied on all listeners’ recordings during an auditory-only probe that was run separately from the main experiment (*Supplementay Method*). The resulting DSS component with the highest evoked activity ratio was used as a single spatial filter for all participants (*Supplementary Fig. 1A*); participants’ DSS time series were downsampled to 32 Hz.

#### 2.5.2 Temporal response function (TRF) estimation

The stimulus time series was indexed by tone pip onsets, a sparse representation of the auditory stimulus *S*(*t*) consisting of a vector equal to 1 at *t*=*T_i_*, the onset time of any given pip, and zero elsewhere. In any AV stimulus, single pips could lead associated flips (auditory precedes visual, ‘ApV’ pips) or lag them (visual precedes auditory, ‘VpA’ pips) with equal probability. Hence the auditory stimuli time series *S*(*t*) was decomposed as *S*(*t*)=*S_VpA_*(*t*)+*S_ApV_*(*t*), where the two time series index the sign of delays between pip onset times *T_i_* and flip times *t_i_*:

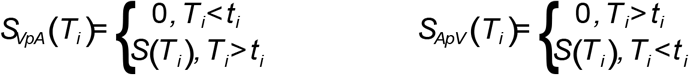

To address the effects of attentional selection, the pips in each of the auditory time series were further decomposed according to the location of associated flips. For instance, in a AH stimulus the time series representing VpA pips was decomposed as *S_VpA.AH_*(*t*)= *S_VpA,AH_ATT__*(*t*)+*S_VpA,AH_UAT__*(*t*). In this case, *AH_ATT_* marks all pips associated with the attended visual hemifield in that presentation, while *AH_UAT_* represents the rest. For AQ stimuli the decomposition was done by quadrants, with equal number of pips per quadrant (see *Supplementary Method*). VpA and ApV defined the *precedence* conditions of pips in a TRF, while AH and AQ stimuli specified their *spatial window* conditions. Finally, two *temporal window* conditions were defined by the AV asynchrony range of the stimulus: stimuli were deemed high AV precision (“Hi-AV”) if their foreground events were all within ±165 ms asynchrony ranges, the remainder were low AV precision “Lo-AV” stimuli (*Supplementary Method*). TRFs were estimated by boosting (David et al., 2007) between concatenated single-trial stimulus and EEG time series, scaled to *z*-units, with 10-fold cross-validation, and assessed at the -70 to 530 ms post-pip onset window. TRFs were additionally estimated from AA stimuli, although these were not used to address the effects of selective attention. Grand average TRFs were obtained by averaging individual TRFs across participants.

### 2.6 Statistical analysis

The experimental conditions of precedence, visuospatial window foreground size, and temporal AV window precision were addressed to: first, localize cross-modal effects on auditory encoding under selective visuospatial attention, and; second, to investigate the interactions between the three conditions on the observed cross-modal effects. For this, in the first step, time intervals indicating differential auditory encoding by visuospatial attention were investigated by cluster-based non-parametric testing (Maris, 2012; Maris & Oostenveld, 2007) corrected for multiple comparisons (see *Supplementary Method*). Upon identification of significant cluster intervals, in the second step their corresponding contrast values (attended minus unattended TRF) were submitted to a repeated measures ANOVA following the factorial structure of the experiment, namely, precedence (VpA, ApV) x foreground size (AH, AQ) x AV precision (Hi-AV, Lo-AV). Post-hoc analyses addressing enhancement effects of attention on individual TRF peak processing were performed with one-sided Wilcoxon signed rank tests at the α=0.05 level of significance, with corrections for multiple comparisons where indicated.

## 3. Results

To determine the presence and timing of modulatory effects of visuospatial selection on auditory processing during AV competition, reproducible auditory neural activity was first extracted from participants by means of data-driven spatial filtering during an auditory-only presentation task. The resulting topography (*Supplementary Fig. 1A*) was consistent with an auditory source (Stropahl et al., 2018) and single-trial EEG data from the two-interval forced choice task were projected to this component, to be submitted to the TRF estimation method. Participants were able to perform the 2-IFC task across conditions (*Supplementary Results*). Pips in the task were partitioned according to their correspondence to: attended versus unattended visual sectors, and pip versus flip precedence (see 2.5.2 *TRF estimation*). TRFs were separately estimated from AH versus AQ, and from Lo- versus Hi-AV precision trials, in both attentional conditions (Figure 3A). We additionally obtained TRFs from AA trials (*Supplementary Fig. 1B*), although these were not used in the attentional analyses. To address the effects of attention, TRF estimates for pips corresponding to the Attended foreground were compared with those associated with the ongoing Unattended background, i.e. opposite hemifield in AH or opposite quadrant in AQ (Figure 3B, C).

**Figure 3.**
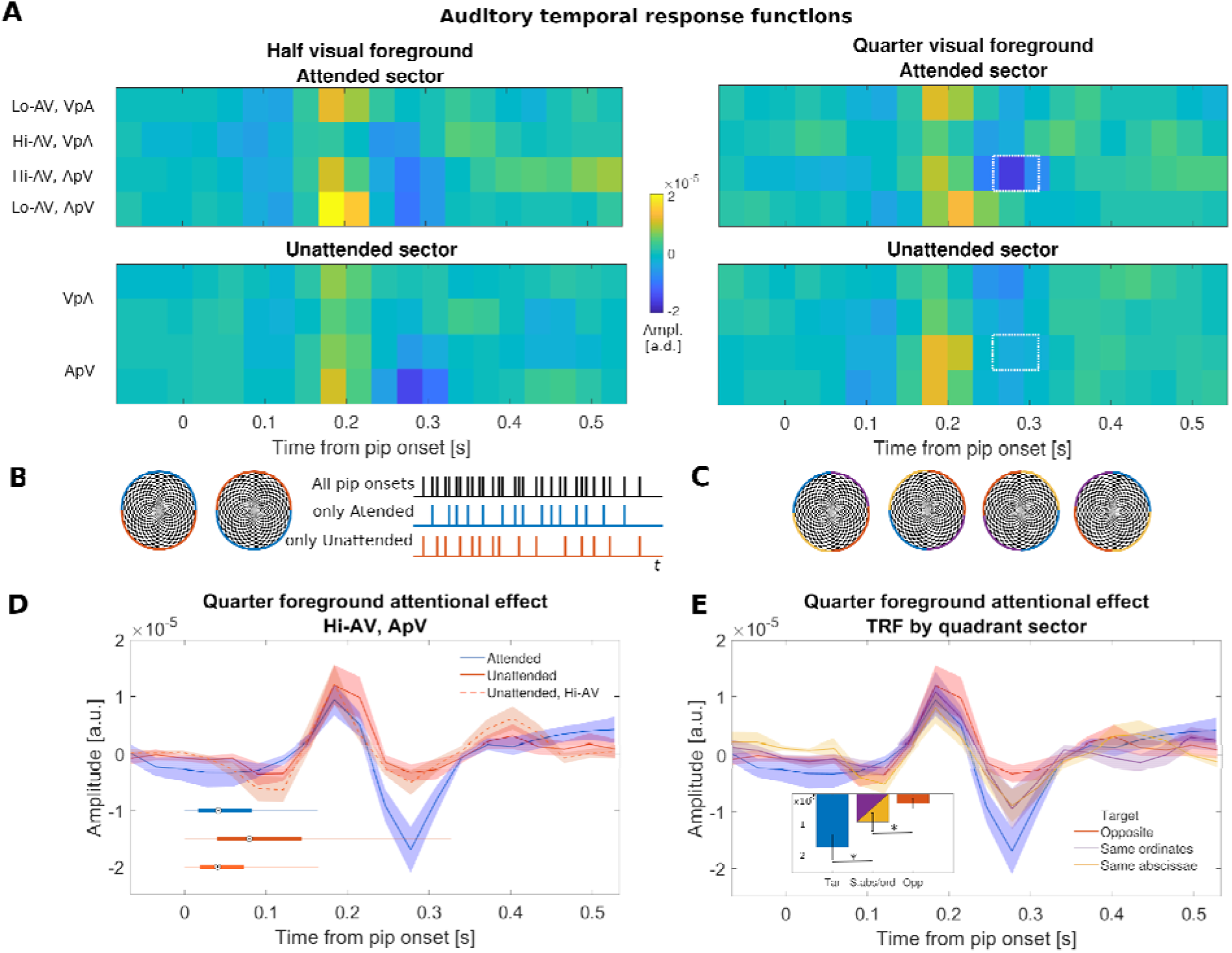
Differential processing of auditory stimuli by visuospatial criteria. (A) Grand average TRFs from Attend-Half (left) and Attend-Quarter trials (right). With AQ foregrounds, neural pip encoding is modulated cross-modally by selective visuospatial attention in the 260 – 320 ms interval (highlighted in white), provided tone onsets closely precede incoming visual input. (B) Illustration of attended sectors in AH trials. Participants attended to visual events from top or bottom hemifields. TRF estimates were obtained from partitioned onset representations of the auditory stream, where only pips associated with the attended or unattended hemifield are represented. (C) In AQ trials, the unattended quadrant opposite to the target (‘Unattended’ in A, D) is flanked by two other unattended quadrants, indicated by yellow and purple here. (D) TRFs where a cross-modal transfer effect was observed, displayed as in A (blue and red) along with corresponding AV onset asynchrony distributions. The pips associated with the Unattended sector entail comparatively lower AV precision; controlling for this parameter (Unattended Hi-AV, orange) similarly results in differential selectivity. (E) Comparison of target quadrant, opposite, and flanking quadrants’ TRFs (blue and red, as in D) shows the flanking quadrant TRFs yield, on average, intermediate levels of representation, consistent with the spatial selection criteria required in the task (inset).

The presence and times at which differential auditory encoding may peak by visuospatial attention were first examined by non-parametric permutation testing corrected for multiple comparisons. The test revealed a significant effect of visuospatial attentional selection on the auditory TRF in the 262-324 ms post pip onset range (*p*<0.001). Greater peak amplitude was observed for pips associated with ‘Attended’ than ‘Unattended’ visual sectors. This effect was observed at Hi-AV AQ trials and for pips closely followed by flips (ApV), and was located in time at the second TRF peak stage of auditory stream processing during AV presentations (Figure 3A, *Supplementary Fig. 1B,C*).

As the permutation analyses first served to localize the presence of a cross-modal transfer attentional effect, the statistical interactions involved between the experimental conditions were next addressed. The Attended minus Unattended TRF contrasts within the identified time period were submitted to a repeated measures 3-way ANOVA with precedence (ApV vs. VpA), temporal (Hi-AV vs. Lo-AV) and spatial window (AH vs. AQ) as factors. There was no significant 3-way interaction (*F*(1,26) = 0.899, *p* = 0.352), and no significant precedence by temporal window interaction (*F*(1,26) = 1.811, *p* = 0.19) or spatial window by temporal window interaction (*F*(1,26) = 0.04, *p* = 0.843). There was, however, a significant precedence by spatial window interaction (*F*(1,26) = 4.643, *p* = 0.041, η^2^=0.152). *Supplementary Fig. 2* presents the data in a form relevant for interpreting the significant interaction. Post-hoc tests show the interaction was due to the lack of a precedence effect, on TRF contrasts, for half (AH) foregrounds (*F*(1, 29) = 0.615, *p* = 0.439); while the effect of precedence was significant for quarter (AQ) foregrounds (F(1,29)□=□5.371; *p*□=□0.028). Finally, the omnibus analysis did not show significant main effects of precedence (*F*(1,26) = 4.903, *p* = 0.717) or spatial window (*F*(1,26) < 0.01, *p* > 0.99), but did reveal a significant main effect of temporal window (*F*(1,26) = 4.903, *p* = 0.036, η^2^=0.159). This effect reflected the fact that TRF contrasts from the pooled Hi-AV conditions were significantly different from zero (attentional effect), while those from Lo-AV conditions were not (two-sided Wilcoxon tests, *p*=0.026 and *p*=0.349 respectively, Bonferroni adjusted). The results overall suggest that cross-modal attentional changes were enabled by visual updating mechanisms under AQ conditions. Since generally only Hi-AV conditions were associated with an attentional modulation, data from Lo-AV temporal window conditions were not further analyzed.

To rule out the possibility that the observed interaction between precedence and spatial window may be due to the relatively longer duration of AQ conditions, data from only the first half of AQ stimuli presentations (i.e. matching AH conditions) were then investigated for TRF estimation. Upon reanalysis, the interaction findings were replicated (*Supplementary Results*). We additionally examined whether AV precision differences at the foreground relative to the background sectors may contribute to the observed effects. Stimuli were constructed in this way to prevent a generic ‘AA’ strategy from participants across trials. To address this, ‘Unattended’ TRFs were estimated again based on all epochs where background pips were also of Hi-AV precision, resulting in similar results (Figure 3D, *Supplementary Results*). The findings confirm that the second TRF peak is subject to enhanced attentional changes by visual updating under conditions of reduced foreground and temporal window size, i.e. lower visuospatial and AV temporal uncertainty. Importantly, the observed auditory changes could not be solely explained by the attentionally biased visual responses which, as expected, showed significant attentional effects both at Half and Quarter foreground sizes (*Supplementary Fig. 3*). This suggests that AH conditions do not efficiently transfer present visual biases onto auditory elements, whereas for AQ this may be the case.

Importantly, AQ transfer dependence on pip precedence was found to be robust to AV associations that may emerge by chance if participants interpret alternative pip onsets as better aligned in time with target flips or viceversa (*Supplementary Results*). Thus, next we investigated the conditions of AV temporal integration in detail. A possibility is that a higher likelihood of onset synchronization explains the AQ results, in partial agreement with transfer by temporal alignment, rather than by temporal integration. To address the impact of temporal proximity between individual pip and flip pairs on the observed cross-modal transfer, AQ ApV pip were separated based on the magnitude of their pip-flip asynchrony by a median split procedure. Here, higher synchrony pips (i.e. onset leads of <39 ms) would be more likely to show cross-modal transfer, according to the temporal matching hypothesis. However, the attentional effects were significant at pips from the *lower* synchrony tier (onset lead range 39 – 165 ms; *p*=7.5 x 10^-4^; *Supplementary Fig. 4*). The result further lends support to the interpretation of effective cross-modal transfer by a visually weighed transient signal onto an auditory process that is already being sustained cortically, i.e. visual updating.

So far, the results identify the stage and spatiotemporal contingencies at which auditory encoding is modulated by visuospatial selection. This neural phenomenon emerges from the interaction between key parameters of AV stimuli: (i) it is conditioned on a limited AV temporal window (<165 ms); (ii) the updating mechanism of transfer over leading sound is contingent on a relatively narrow target area (1:4 foreground to background ratio). One may then hypothesize that, upon transfer, the emergent auditory representation being selected may inherit spatial-like attributes. We next investigated the representation of unattended pips at areas that are spatially closer to the target, i.e. flanking quadrants. Pip encoding of the two unattended quadrant sectors between the target and its opposite quadrant – i.e., same abscissae and same ordinates quadrants (Figure 3C) was addressed. TRF estimates from pips associated with these sectors illustrate how, for such cases, the second TRF peak encodes pips at intermediate amplitudes between target and opposite quadrant levels (Figure 3E). A one-sided Wilcoxon signed-rank test of the average TRF amplitude across both flanking quadrants showed a significantly greater TRF peak amplitude for pips on target (Figure 3E, D, blue) than pips on flanking quadrants (Figure 3E, yellow and purple) (*Z*=-2.03, adjusted *p*=0.042, Bonferroni corrected), in the 262 – 293 ms window. Additionally, the average TRF peak amplitude for pips on flanking quadrants was significantly greater than that of the quadrant opposite to the target (Figure 3E, D, red) (*Z*=-2.13, adjusted *p*=0.034), for the same temporal window.

## 4. Discussion

How do visuospatial biases transfer effectively into the neural encoding of auditory streams has remained, to our knowledge, an unaddressed question so far. The present study considered key distributional properties of AV stimulation that may account for optimal cross-modal transfer. Participants monitored selected visual locations to determine the degree of asynchrony between probabilistic visual and auditory sequences. Changes to their auditory encoding during the task were investigated by means of EEG reverse correlation techniques. Evidence of cross-modal transfer was observed at 300 ms post audio onset, and was contingent on auditory-over-visual lead, narrow AV asynchrony margins (~39 ms), and a limited (1:4) visual foreground to background ratio. The results demonstrate that neural auditory segregation in competitive task scenarios can be directed visuospatially. The dependence of the transfer effect on the arrival of visual transients during, or shortly after, leading sound presentations underscores the visual updating of ongoing auditory processes. The findings also suggest that auditory segregation in the task was based on pips’ acquisition of spatial information that was originally available from the visual stream. Such changes to the encoding of sound are consistent with the dynamic re-weighting of auditory representations, according to the temporal and spatial proximity to a cross-modally selected region.

Rather than an automatic type of AV integration (e.g. Koelewijn et al., 2010; Giard & Peronnet, 1999; Kayser et al., 2007; Noesselt et al., 2007), the transfer’s relatively late timing and its dependence on auditory over visual precedence support the interpretation of modified pip representations as they become part of multimodal AV objects with visually attended features. For comparison, in vision, unselected spatial locations that happen to be part of a foreground object may undergo enhanced processing after 180 ms of visual stimulus onset (Martínez et al., 2006). This phenomenon of transfer, across independent feature dimensions of an object, is defined as “binding” and also applies to multimodal scenarios (Bizley et al., 2016; Donohue et al., 2011; Spence & Frings, 2020; Talsma et al., 2010). Clearly, both unimodal and bimodal forms of binding involve contextual or high-level object processing, while bimodal cases additionally require specific associative processes that are not modality-specific. Attention, for instance, may precede certain forms of cross-modal integration (Ernst & Bülthoff, 2004; Rohe et al., 2019; Spence & Ngo, 2012). It has been noted that AV integration processes may initiate outside the scope of attention (Atilgan et al., 2018), yet the balance of attentional deployment is known to affect multimodal object representations (Badde et al., 2020; French & DeAngelis, 2020; Rohe et al., 2019). With hearing’s temporal processing advantages, sound is well suited to rapidly signal observers to engage visual attention (e.g. Spence & Driver, 1997; Fujisaki et al., 2004). Leading sound stimuli often mark when to enhance attentional deployment at specified visual locations, and to integrate visual feedback thereafter (Noesselt et al., 2008; Shams et al., 2005; Van der Burg et al., 2008, 2011). The present results showed that at sound-leading conditions auditory processing may be modulated visuospatially. These represent the greatest chance for individual visual contrast changes to coincide with the sustained phase of individual sound presentations, when auditory processes may already take place cortically. Thus, our findings support the role of visual updating mechanisms in auditory scene analysis under competitive AV selection using visuospatial criteria (Koelewijn et al., 2009; Roseboom et al., 2009). Future studies may conversely address the role of auditory updating on visual representations under unimodal auditory selection.

The present results further illustrate how multimodal stimuli do not need to be simultaneous to be inferred as being produced from a single source, as in naturalistic scenarios oftentimes. Robust AV object perception flexibly allows for a certain degree of temporal misalignment (van Wassenhove et al., 2007). At the neural coding level, our results accordingly suggest that cross-modal transfer is still possible under relatively uncertain temporal AV associations. Notably, transfer was also contingent on constraints to spatial foreground size. One possible explanation for this could be that AH pips may be individually less effective at tagging visual processing for enhanced attention (e.g., Olivers & Van der Burg, 2008; Quak et al., 2015). Alternatively, AQ window spans facilitated transfer because they distribute the total visual perceptual load (Lavie, 1995) over fewer spatial channels, hence optimizing serial deployments of visuospatial attention unimodally (Fujisaki et al., 2006; Andersen et al., 2009). Two present results support this latter interpretation, first, that segregated auditory representations were driven by inferred spatial proximity to the visual target via temporal association. Second, our control TRF analyses for the AV precision of background pips showed that attentional segregation on a visuospatial basis was possible regardless of higher AV precision in the background area; moreover, it was not determined by the particular dynamics of the auditory background during competition (cf. Rahne et al., 2007).

In the last decade, several computational frameworks of attention have proposed that perceptual systems actively estimate noise in order to weigh sensory data for selection (Hesselmann et al., 2010; Macaluso et al., 2016; Rohe & Noppeney, 2018; Yu, 2014). In these models, the brain discounts representations of sensory input according to representations of their inferred uncertainty. After normalizing sensory input by their relative uncertainty, attention could operate by selecting the inputs with resulting higher signal-to-noise ratio (Yu & Dayan, 2002, 2005b). Therefore to influence a neural representation, one may externally change the input itself or alternatively its associated perceptual uncertainty. The term ‘uncertainty’ is concerned with the reliability of external sensory variables for perceptual inference; this refers to the principle by which observers generally predict the sort of sensory input they will encounter throughout their world – observers continually assess their confidence in their supplied sensory data given what they know about how those same data are generated (Parr & Friston, 2019). In the task of this study, judgment of temporal variance was actively required from participants. Consequently, the findings showed that attentional modulations not only depended on bimodal uncertainty (asynchrony) but also unimodal uncertainty (position) terms. Reducing these uncertainties in the stimulus and the target space had the combined effect of *updating* those tone representations that potentially match as segregated elements of an attended multimodal object (cf. Beauchamp et al., 2004; Noesselt et al., 2007; Talsma et al., 2010). By parameterizing the likelihood of a match, or its quality, AV temporal uncertainty appears suited to influence both attentional selection and cross-modal binding. Interestingly, both aspects have been shown to operate on partially overlapping circuitry throughout the depth axis of the auditory cortex (Gau et al., 2020). In addition, sampling visual target input with higher positional uncertainty constrained the ability for visuospatial biases to be transferred cross-modally. There is evidence that uncertainty representation in unisensory cortex is relevant for behavior (Walker et al., 2020), and that unimodal transfer of selective biases across visual features may be facilitated by reductions to unisensory uncertainty (Luo et al., 2018). Here, AQ would rely on comparatively fewer sensory neurons than at AH, where a greater proportion of visual sensory inputs should not be attenuated, inhibited, or discarded. Thus by indexing fewer locations, relative visual weights could bind more effectively into auditory selective processes. Other computational strategies that account for sensory uncertainty during fused audiovisual perception have been reported in the cortex (Rohe et al., 2019; Rohe & Noppeney, 2015; Meijer et al., 2019).

## 5. Conclusion

Optimal attentional selection relies on adopting statistical properties of natural signals that help enhance their representation relative to distractors, according to task demands. In multisensory integration, a key property relates to the quality of the temporal association between unimodal streams. The present data confirm that such temporal factor determines whether top-down visuospatial biases transmit cross-modally, in the process of updating the neural representations of ongoing sound. Another key factor involves the unimodal uncertainty associated with visual foreground top-down selection. Under visuospatial attention, reducing foreground size may promote the re-weighting of the neural representation of sounds by their inferred proximity to target criteria, in effect segregating relevant elements of the auditory stream.

## Author Notes

We thank Michelle Symonds for English editing. The research presented in this manuscript received funding from the Agencia Nacional de Investigación e Innovación, Uruguay, under the code PD_NAC_2018_1_150365.

## Data availability statement

The data supporting the findings of this study are openly available at the Open Science Framework at http://doi.org/10.17605/OSF.IO/8V9SD

---

**Video 1.** Sample Attend-All trial on the 2-IFC task. After presenting the initial cue to attend the full foreground, the two AV sequences are presented in series. In this example, the first stimulus has lower AV temporal precision than the second. On AA trials, the participant is asked to consider all pip-flip pairings across each of the two disc presentations. After the second presentation, participants decide which of the two sequences had comparatively less (or more) temporal precision. Feedback is shown upon response validation. An example selection of the correct response is also represented here.

## References

Andersen, T. S., Tiippana, K., Laarni, J., Kojo, I., & Sams, M. (2009). The role of visual spatial attention in audiovisual speech perception. Speech Communication, 51(2), 184–193. https://doi.org/10.1016/j.specom.2008.07.004

Atilgan, H., Town, S. M., Wood, K. C., Jones, G. P., Maddox, R. K., Lee, A. K. C., & Bizley, J. K. (2018). Integration of Visual Information in Auditory Cortex Promotes Auditory Scene Analysis through Multisensory Binding. Neuron, 97(3), 640–655.e4. https://doi.org/10.1016/j.neuron.2017.12.034

Badde, S., Navarro, K. T., & Landy, M. S. (2020). Modality-specific attention attenuates visual-tactile integration and recalibration effects by reducing prior expectations of a common source for vision and touch. Cognition, 197, 104170. https://doi.org/10.1016/j.cognition.2019.104170

Beauchamp, M. S. (2019). Using Multisensory Integration to Understand the Human Auditory Cortex. In A. K. C. Lee, M. T. Wallace, A. B. Coffin, A. N. Popper, & R. R. Fay (Eds.), Multisensory Processes: The Auditory Perspective (pp. 161–176). Springer International Publishing. https://doi.org/10.1007/978-3-030-10461-0_8

Beauchamp, M. S., Lee, K. E., Argall, B. D., & Martin, A. (2004). Integration of Auditory and Visual Information about Objects in Superior Temporal Sulcus. Neuron, 41(5), 809–823. https://doi.org/10.1016/S0896-6273(04)00070-4

Bizley, J. K., Maddox, R. K., & Lee, A. K. C. (2016). Defining Auditory-Visual Objects: Behavioral Tests and Physiological Mechanisms. Trends in Neurosciences, 39(2), 74–85. https://doi.org/10.1016/j.tins.2015.12.007

Brodbeck, C., Jiao, A., Hong, L. E., & Simon, J. Z. (2020). Neural speech restoration at the cocktail party: Auditory cortex recovers masked speech of both attended and ignored speakers. PLOS Biology, 18(10), e3000883. https://doi.org/10.1371/journal.pbio.3000883

Burr, D., Banks, M. S., & Morrone, M. C. (2009). Auditory dominance over vision in the perception of interval duration. Experimental Brain Research, 198(1), 49–57. doi: 10.1007/s00221-009-1933-z

Busse, L., Katzner, S., & Treue, S. (2008). Temporal dynamics of neuronal modulation during exogenous and endogenous shifts of visual attention in macaque area MT. Proceedings of the National Academy of Sciences of the United States of America, 105(42), 16380–16385. https://doi.org/10.1073/pnas.0707369105

Capilla, A., Melcón, M., Kessel, D., Calderón, R., Pazo-Álvarez, P., & Carretié, L. (2016). Retinotopic mapping of visual event-related potentials. Biological Psychology, 118, 114–125. https://doi.org/10.1016/j.biopsycho.2016.05.009

Carrasco, M. (2014). Spatial Covert Attention. The Oxford Handbook of Attention. https://doi.org/10.1093/oxfordhb/9780199675111.013.004

Cervantes Constantino, F., Villafañe-Delgado, M., Camenga, E., Dombrowski, K., Walsh, B., & Simon, J. Z. (2017). Functional significance of spectrotemporal response functions obtained using magnetoencephalography. BioRxiv, 168997. https://doi.org/10.1101/168997

Crosse, M. J., Di Liberto, G. M., Bednar, A., & Lalor, E. C. (2016). The Multivariate Temporal Response Function (mTRF) Toolbox: A MATLAB Toolbox for Relating Neural Signals to Continuous Stimuli. Frontiers in Human Neuroscience, 10. https://doi.org/10.3389/fnhum.2016.00604

David, S. V., Mesgarani, N., & Shamma, S. A. (2007). Estimating sparse spectro-temporal receptive fields with natural stimuli. Network: Computation in Neural Systems, 18(3), 191–212. https://doi.org/10.1080/09548980701609235

Dayan, P., Kakade, S., & Montague, P. R. (2000). Learning and selective attention. Nature Neuroscience, 3 Suppl, 1218–1223. https://doi.org/10.1038/81504

Dayan, P., & Yu, A. J. (2003). Uncertainty and Learning. IETE Journal of Research, 49(2–3), 171–181. https://doi.org/10.1080/03772063.2003.11416335

Dayan, P., & Zemel, R. S. (1999). Statistical models and sensory attention. 1999 Ninth International Conference on Artificial Neural Networks ICANN 99. (Conf. Publ. No. 470), 2, 1017–1022 vol.2. https://doi.org/10.1049/cp:19991246

de Cheveigné, A., & Parra, L. C. (2014). Joint decorrelation, a versatile tool for multichannel data analysis. NeuroImage, 98, 487–505. https://doi.org/10.1016/j.neuroimage.2014.05.068

de Cheveigné, A., & Simon, J. Z. (2007). Denoising based on time-shift PCA. Journal of Neuroscience Methods, 165(2), 297–305. https://doi.org/10.1016/j.jneumeth.2007.06.003

de Cheveigné, A., & Simon, J. Z. (2008a). Sensor noise suppression. Journal of Neuroscience Methods, 168(1), 195–202. https://doi.org/10.1016/j.jneumeth.2007.09.012

de Cheveigné, A., & Simon, J. Z. (2008b). Denoising based on spatial filtering. Journal of Neuroscience Methods, 171(2), 331–339. https://doi.org/10.1016/j.jneumeth.2008.03.015

Degerman, A., Rinne, T., Pekkola, J., Autti, T., Jääskeläinen, I. P., Sams, M., & Alho, K. (2007). Human brain activity associated with audiovisual perception and attention. NeuroImage, 34(4), 1683–1691. https://doi.org/10.1016/j.neuroimage.2006.11.019

Ding, N., & Simon, J. Z. (2012). Emergence of neural encoding of auditory objects while listening to competing speakers. Proceedings of the National Academy of Sciences, 109(29), 11854–11859. https://doi.org/10.1073/pnas.1205381109

Donohue, S. E., Roberts, K. C., Grent-’t-Jong, T., & Woldorff, M. G. (2011). The Cross-Modal Spread of Attention Reveals Differential Constraints for the Temporal and Spatial Linking of Visual and Auditory Stimulus Events. Journal of Neuroscience, 31(22), 7982–7990. https://doi.org/10.1523/JNEUROSCI.5298-10.2011

Egly, R., Driver, J., & Rafal, R. (1994). Shifting Visual-Attention Between Objects and Locations—Evidence from Normal and Parietal Lesion Subjects. Journal of Experimental Psychology-General, 123(2), 161–177. https://doi.org/10.1037//0096-3445.123.2.161

Eimer, M. (2000). An ERP study of sustained spatial attention to stimulus eccentricity. Biological Psychology, 52(3), 205–220. https://doi.org/10.1016/S0301-0511(00)00028-4

Ernst, M. O., & Bülthoff, H. H. (2004). Merging the senses into a robust percept. Trends in Cognitive Sciences, 8(4), 162–169. https://doi.org/10.1016/j.tics.2004.02.002

Feldman, H., & Friston, K. (2010). Attention, Uncertainty, and Free-Energy. Frontiers in Human Neuroscience, 4. https://doi.org/10.3389/fnhum.2010.00215

Fleming, J. T., Noyce, A. L., & Shinn-Cunningham, B. G. (2020). Audio-visual spatial alignment improves integration in the presence of a competing audio-visual stimulus. Neuropsychologia, 146, 107530. https://doi.org/10.1016/j.neuropsychologia.2020.107530

French, R. L., & DeAngelis, G. C. (2020). Multisensory neural processing: From cue integration to causal inference. Current Opinion in Physiology, 16, 8–13. https://doi.org/10.1016/j.cophys.2020.04.004

Fujisaki, W., Koene, A., Arnold, D., Johnston, A., & Nishida, S. (2006). Visual search for a target changing in synchrony with an auditory signal. Proceedings. Biological Sciences, 273(1588), 865–874. https://doi.org/10.1098/rspb.2005.3327

Fujisaki, W., Shimojo, S., Kashino, M., & Nishida, S. (2004). Recalibration of audiovisual simultaneity. Nature Neuroscience, 7(7), 773–778. https://doi.org/10.1038/nn1268

Gau, R., Bazin, P.-L., Trampel, R., Turner, R., & Noppeney, U. (2020). Resolving multisensory and attentional influences across cortical depth in sensory cortices. ELife, 9, e46856. https://doi.org/10.7554/eLife.46856

Gaucher, Q., Edeline, J.-M., & Gourévitch, B. (2012). How different are the local field potentials and spiking activities? Insights from multi-electrodes arrays. Journal of Physiology-Paris, 106(3–4), 93–103. https://doi.org/10.1016/j.jphysparis.2011.09.006

Giard, M. H., & Peronnet, F. (1999). Auditory-visual integration during multimodal object recognition in humans: A behavioral and electrophysiological study. Journal of Cognitive Neuroscience, 11(5), 473–490. https://doi.org/10.1162/089892999563544

Glasberg, B. R., & Moore, B. C. J. (1990). Derivation of auditory filter shapes from notched-noise data. Hearing Research, 47(1–2), 103–138. https://doi.org/10.1016/0378-5955(90)90170-T

Gourévitch, B., Noreña, A., Shaw, G., & Eggermont, J. J. (2009). Spectrotemporal receptive fields in anesthetized cat primary auditory cortex are context dependent. Cerebral Cortex (New York, N.Y.: 1991), 19(6), 1448–1461. https://doi.org/10.1093/cercor/bhn184

Groen, I. I. A., Dekker, T. M., Knapen, T., & Silson, E. H. (2021). Visuospatial coding as ubiquitous scaffolding for human cognition. Trends in Cognitive Sciences, 0(0). https://doi.org/10.1016/j.tics.2021.10.011

Herrmann, C. S., & Knight, R. T. (2001). Mechanisms of human attention: Event-related potentials and oscillations. Neuroscience & Biobehavioral Reviews, 25(6), 465–476. https://doi.org/10.1016/S0149-7634(01)00027-6

Hesselmann, G., Sadaghiani, S., Friston, K. J., & Kleinschmidt, A. (2010). Predictive coding or evidence accumulation? False inference and neuronal fluctuations. PloS One, 5(3), e9926. https://doi.org/10.1371/journal.pone.0009926

Hillyard, S. A., Hink, R. F., Schwent, V. L., & Picton, T. W. (1973). Electrical signs of selective attention in the human brain. Science (New York, N.Y.), 182(4108), 177–180.

Hollingworth, A., Maxcey-Richard, A. M., & Vecera, S. P. (2012). The spatial distribution of attention within and across objects. Journal of Experimental Psychology. Human Perception and Performance, 38(1), 135–151. https://doi.org/10.1037/a0024463

Holmes, N. P., & Spence, C. (2005). Multisensory integration: Space, time, & superadditivity. Current Biology□: CB, 15(18), R762–R764. https://doi.org/10.1016/j.cub.2005.08.058

Hyvarinen, A. (1999). Fast and robust fixed-point algorithms for independent component analysis. IEEE Transactions on Neural Networks, 10(3), 626–634. https://doi.org/10.1109/72.761722

Jensen, A., Merz, S., Spence, C., & Frings, C. (2019). Overt spatial attention modulates multisensory selection. Journal of Experimental Psychology. Human Perception and Performance, 45(2), 174–188. https://doi.org/10.1037/xhp0000595

Junghöfer, M., Elbert, T., Tucker, D. M., & Rockstroh, B. (2000). Statistical control of artifacts in dense array EEG/MEG studies. Psychophysiology, 37(04), 523–532. https://doi.org/null

Kayser, C., Petkov, C. I., Augath, M., & Logothetis, N. K. (2007). Functional imaging reveals visual modulation of specific fields in auditory cortex. The Journal of Neuroscience: The Official Journal of the Society for Neuroscience, 27(8), 1824–1835. https://doi.org/10.1523/JNEUROSCI.4737-06.2007

King, A. J., Hammond-Kenny, A., & Nodal, F. R. (2019). Multisensory Processing in the Auditory Cortex. In A. K. C. Lee, M. T. Wallace, A. B. Coffin, A. N. Popper, & R. R. Fay (Eds.), Multisensory Processes: The Auditory Perspective (pp. 105–133). Springer International Publishing. https://doi.org/10.1007/978-3-030-10461-0_6

Koelewijn, T., Bronkhorst, A., & Theeuwes, J. (2009). Auditory and visual capture during focused visual attention. Journal of Experimental Psychology. Human Perception and Performance, 35(5), 1303–1315. https://doi.org/10.1037/a0013901

Koelewijn, T., Bronkhorst, A., & Theeuwes, J. (2010). Attention and the multiple stages of multisensory integration: A review of audiovisual studies. Acta Psychologica, 134(3), 372–384. https://doi.org/10.1016/j.actpsy.2010.03.010

Lavie, N. (1995). Perceptual load as a necessary condition for selective attention. Journal of Experimental Psychology. Human Perception and Performance, 21(3), 451–468. https://doi.org/10.1037//0096-1523.21.3.451

Luo, T., Wu, X., Wang, H., & Fu, S. (2018). Prioritization to visual objects: Roles of sensory uncertainty. Attention, Perception, & Psychophysics, 80(2), 512–526. https://doi.org/10.3758/s13414-017-1452-0

Macaluso, E., Noppeney, U., Talsma, D., Vercillo, T., Hartcher-O’Brien, J., & Adam, R. (2016). The Curious Incident of Attention in Multisensory Integration: Bottom-up vs. Top-down. Multisensory Research, 29(6–7), 557–583. https://doi.org/10.1163/22134808-00002528

Maddox, R. K., Atilgan, H., Bizley, J. K., & Lee, A. K. C. (2015). Auditory selective attention is enhanced by a task-irrelevant temporally coherent visual stimulus in human listeners. ELife, 4. https://doi.org/10.7554/eLife.04995

Mäkelä, A. M., Alku, P., Mäkinen, V., Valtonen, J., May, P., & Tiitinen, H. (2002). Human cortical dynamics determined by speech fundamental frequency. NeuroImage, 17(3), 1300–1305.

Maris, E. (2012). Statistical testing in electrophysiological studies. Psychophysiology, 49(4), 549–565. https://doi.org/10.1111/j.1469-8986.2011.01320.x

Maris, E., & Oostenveld, R. (2007). Nonparametric statistical testing of EEG- and MEG-data. Journal of Neuroscience Methods, 164(1), 177–190. https://doi.org/10.1016/j.jneumeth.2007.03.024

Martínez, A., Teder-Sälejärvi, W., Vazquez, M., Molholm, S., Foxe, J. J., Javitt, D. C., Di Russo, F., Worden, M. S., & Hillyard, S. A. (2006). Objects Are Highlighted by Spatial Attention. Journal of Cognitive Neuroscience, 18(2), 298–310. https://doi.org/10.1162/jocn.2006.18.2.298

McGurk, H., & MacDonald, J. (1976). Hearing lips and seeing voices. Nature, 264(5588), 746–748. https://doi.org/10.1038/264746a0

Meijer, D., Veselič, S., Calafiore, C., & Noppeney, U. (2019). Integration of audiovisual spatial signals is not consistent with maximum likelihood estimation. Cortex; a Journal Devoted to the Study of the Nervous System and Behavior, 119, 74–88. https://doi.org/10.1016/j.cortex.2019.03.026

Miller, L. M., & D’Esposito, M. (2005). Perceptual fusion and stimulus coincidence in the cross-modal integration of speech. The Journal of Neuroscience: The Official Journal of the Society for Neuroscience, 25(25), 5884–5893. https://doi.org/10.1523/JNEUROSCI.0896-05.2005

Müller, N. G., & Kleinschmidt, A. (2003). Dynamic Interaction of Object- and Space-Based Attention in Retinotopic Visual Areas. Journal of Neuroscience, 23(30), 9812–9816. https://doi.org/10.1523/JNEUROSCI.23-30-09812.2003

Noesselt, T., Bergmann, D., Hake, M., Heinze, H.-J., & Fendrich, R. (2008). Sound increases the saliency of visual events. Brain Research, 1220, 157–163. https://doi.org/10.1016/j.brainres.2007.12.060

Noesselt, T., Fendrich, R., Bonath, B., Tyll, S., & Heinze, H.-J. (2005). Closer in time when farther in space—Spatial factors in audiovisual temporal integration. Cognitive Brain Research, 25(2), 443–458. doi: 10.1016/j.cogbrainres.2005.07.005

Noesselt, T., Rieger, J. W., Schoenfeld, M. A., Kanowski, M., Hinrichs, H., Heinze, H.-J., & Driver, J. (2007). Audiovisual temporal correspondence modulates human multisensory superior temporal sulcus plus primary sensory cortices. The Journal of Neuroscience: The Official Journal of the Society for Neuroscience, 27(42), 11431–11441. https://doi.org/10.1523/JNEUROSCI.2252-07.2007

O’Connor, D. H., Fukui, M. M., Pinsk, M. A., & Kastner, S. (2002). Attention modulates responses in the human lateral geniculate nucleus. Nature Neuroscience, 5(11), 1203–1209. https://doi.org/10.1038/nn957

Olivers, C. N. L., & Van der Burg, E. (2008). Bleeping you out of the blink: Sound saves vision from oblivion. Brain Research, 1242, 191–199. https://doi.org/10.1016/j.brainres.2008.01.070

O’Sullivan, J. A., Power, A. J., Mesgarani, N., Rajaram, S., Foxe, J. J., Shinn-Cunningham, B. G., Slaney, M., Shamma, S. A., & Lalor, E. C. (2015). Attentional Selection in a Cocktail Party Environment Can Be Decoded from Single-Trial EEG. Cerebral Cortex, 25(7), 1697–1706. https://doi.org/10.1093/cercor/bht355

Parr, T., & Friston, K. J. (2019). Attention or salience? Current Opinion in Psychology, 29, 1–5. https://doi.org/10.1016/j.copsyc.2018.10.006

Peirce, J. W. (2007). PsychoPy—Psychophysics software in Python. Journal of Neuroscience Methods, 162(1), 8–13. https://doi.org/10.1016/j.jneumeth.2006.11.017

Quak, M., London, R. E., & Talsma, D. (2015). A multisensory perspective of working memory. Frontiers in Human Neuroscience, 9. https://doi.org/10.3389/fnhum.2015.00197

Rahne, T., Böckmann, M., von Specht, H., & Sussman, E. S. (2007). Visual cues can modulate integration and segregation of objects in auditory scene analysis. Brain Research, 1144, 127–135. https://doi.org/10.1016/j.brainres.2007.01.074

Roberts, T. P., Ferrari, P., Stufflebeam, S. M., & Poeppel, D. (2000). Latency of the auditory evoked neuromagnetic field components: Stimulus dependence and insights toward perception. Journal of Clinical Neurophysiology: Official Publication of the American Electroencephalographic Society, 17(2), 114–129. https://doi.org/10.1097/00004691-200003000-00002

Rohe, T., Ehlis, A.-C., & Noppeney, U. (2019). The neural dynamics of hierarchical Bayesian causal inference in multisensory perception. Nature Communications, 10(1), 1–17. https://doi.org/10.1038/s41467-019-09664-2

Rohe, T., & Noppeney, U. (2015). Cortical Hierarchies Perform Bayesian Causal Inference in Multisensory Perception. PLOS Biology, 13(2), e1002073. https://doi.org/10.1371/journal.pbio.1002073

Rohe, T., & Noppeney, U. (2018). Reliability-Weighted Integration of Audiovisual Signals Can Be Modulated by Top-down Attention. ENeuro, 5(1). https://doi.org/10.1523/ENEURO.0315-17.2018

Roseboom, W., Nishida, S., & Arnold, D. H. (2009). The sliding window of audio-visual simultaneity. Journal of Vision, 9(12), 4.1–8. https://doi.org/10.1167/9.12.4

Santangelo, V. & Macaluso, E. (2012). Spatial attention and audiovisual processing. In The New Handbook of Multisensory Processing. MIT Press. https://www.academia.edu/16745204/Spatial_attention_and_audiovisual_processing

Shams, L., Iwaki, S., Chawla, A., & Bhattacharya, J. (2005). Early modulation of visual cortex by sound: An MEG study. Neuroscience Letters, 378(2), 76–81. https://doi.org/10.1016/j.neulet.2004.12.035

Shinn-Cunningham, B. G. (2008). Object-based auditory and visual attention. Trends in Cognitive Sciences, 12(5), 182–186. https://doi.org/10.1016/j.tics.2008.02.003

Spence, C., & Driver, J. (1997). Audiovisual links in exogenous covert spatial orienting. Perception & Psychophysics, 59(1), 1–22. https://doi.org/10.3758/bf03206843

Spence, C., & Frings, C. (2020). Multisensory feature integration in (and out) of the focus of spatial attention. Attention, Perception, & Psychophysics, 82(1), 363–376. https://doi.org/10.3758/s13414-019-01813-5

Spence, C., & Ngo, M. K. (2012). Does Attention or Multisensory Integration Explain the Cross-Modal Facilitation of Masked Visual Target Identification? In Stein Barry E. (Ed.), The New Handbook of Multisensory Processing (1st edition). The MIT Press. http://cognet.mit.edu/erefs/new-handbook-of-multisensory-processing

Spence, C., & Squire, S. (2003). Multisensory Integration: Maintaining the Perception of Synchrony. Current Biology, 13(13), R519–R521. https://doi.org/10.1016/S0960-9822(03)00445-7

Stropahl, M., Bauer, A.-K. R., Debener, S., & Bleichner, M. G. (2018). Source-Modeling Auditory Processes of EEG Data Using EEGLAB and Brainstorm. Frontiers in Neuroscience, 12. https://doi.org/10.3389/fnins.2018.00309

Talsma, D., Senkowski, D., Soto-Faraco, S., & Woldorff, M. G. (2010). The multifaceted interplay between attention and multisensory integration. Trends in Cognitive Sciences, 14(9), 400–410. https://doi.org/10.1016/j.tics.2010.06.008

Talsma, D., & Woldorff, M. G. (2005). Selective attention and multisensory integration: Multiple phases of effects on the evoked brain activity. Journal of Cognitive Neuroscience, 17(7), 1098–1114. https://doi.org/10.1162/0898929054475172

Treisman, A. (1998). Feature binding, attention and object perception. Philosophical Transactions of the Royal Society B: Biological Sciences, 353(1373), 1295–1306.

Treisman, A. M., & Gelade, G. (1980). A feature-integration theory of attention. Cognitive Psychology, 12(1), 97–136. https://doi.org/10.1016/0010-0285(80)90005-5

van Atteveldt, N. M., Formisano, E., Blomert, L., & Goebel, R. (2007). The effect of temporal asynchrony on the multisensory integration of letters and speech sounds. Cerebral Cortex (New York, N.Y.: 1991), 17(4), 962–974. https://doi.org/10.1093/cercor/bhl007

Van der Burg, E., Cass, J., Olivers, C. N. L., Theeuwes, J., & Alais, D. (2010). Efficient Visual Search from Synchronized Auditory Signals Requires Transient Audiovisual Events. PLOS ONE, 5(5), e10664. https://doi.org/10.1371/journal.pone.0010664

Van der Burg, E., Olivers, C. N. L., Bronkhorst, A. W., & Theeuwes, J. (2008). Pip and pop: Nonspatial auditory signals improve spatial visual search. Journal of Experimental Psychology. Human Perception and Performance, 34(5), 1053–1065. https://doi.org/10.1037/0096-1523.34.5.1053

Van der Burg, E., Talsma, D., Olivers, C. N. L., Hickey, C., & Theeuwes, J. (2011). Early multisensory interactions affect the competition among multiple visual objects. NeuroImage, 55(3), 1208–1218. https://doi.org/10.1016/j.neuroimage.2010.12.068

van Wassenhove, V., Grant, K. W., & Poeppel, D. (2007). Temporal window of integration in auditory-visual speech perception. Neuropsychologia, 45(3), 598–607. https://doi.org/10.1016/j.neuropsychologia.2006.01.001

Walker, E. Y., Cotton, R. J., Ma, W. J., & Tolias, A. S. (2020). A neural basis of probabilistic computation in visual cortex. Nature Neuroscience, 23(1), 122–129. https://doi.org/10.1038/s41593-019-0554-5

Wannig, A., Stanisor, L., & Roelfsema, P. R. (2011). Automatic spread of attentional response modulation along Gestalt criteria in primary visual cortex. Nature Neuroscience, 14(10), 1243–1244. https://doi.org/10.1038/nn.2910

Whiteley, L., & Sahani, M. (2008). Implicit knowledge of visual uncertainty guides decisions with asymmetric outcomes. Journal of Vision, 8(3), 2.1–215. https://doi.org/10.1167/8.3.2

Whiteley, L., & Sahani, M. (2012). Attention in a Bayesian Framework. Frontiers in Human Neuroscience, 6. https://doi.org/10.3389/fnhum.2012.00100

World Medical Association (WMA). (2009). Declaration of Helsinki. Ethical Principles for Medical Research Involving Human Subjects. Jahrbuch Für Wissenschaft Und Ethik, 14(1), 233–238. https://doi.org/10.1515/9783110208856.233

Yu, A. J. (2014). Bayesian Models of Attention. In A. C. Nobre & S. Kastner (Eds.), The Oxford Handbook of Attention. Oxford University Press. https://doi.org/10.1093/oxfordhb/9780199675111.013.025

Yu, A. J., & Dayan, P. (2002). Acetylcholine in cortical inference. Neural Networks: The Official Journal of the International Neural Network Society, 15(4–6), 719–730. https://doi.org/10.1016/s0893-6080(02)00058-8

Yu, A. J., & Dayan, P. (2005a). Inference, Attention, and Decision in a Bayesian Neural Architecture. In L. K. Saul, Y. Weiss, & L. Bottou (Eds.), Advances in Neural Information Processing Systems 17 (pp. 1577–1584). MIT Press. http://papers.nips.cc/paper/2548-inference-attention-and-decision-in-a-bayesian-neural-architecture.pdf

Yu, A. J., & Dayan, P. (2005b). Uncertainty, neuromodulation, and attention. Neuron, 46(4), 681–692. https://doi.org/10.1016/j.neuron.2005.04.026

