## Supplementary Material for "Visuospatial attention revamps cortical processing of sound: restrict stimulus uncertainty"

**Supplementary Method**

**Stimuli**

*Audiovisual stimuli*. Asynchronies between consecutive flips were defined as 101 ms plus an exponentially distributed random delay (μ = 99 ms). This nominal asynchrony was subsequently discretized for correspondence with the video file frame rate (10 fps) by increments of 100 ms. A frequency-eccentricity association was introduced so that the lowest three pip frequencies (100, 171, 257 Hz) were reserved for flips located at the most peripheral eccentricity. Likewise, the highest three frequencies (3.2, 3.9, 4.8 KHz) were used only for flips at the most central eccentricity, while intermediate frequencies were arranged accordingly. These mappings were intended to counter, on the auditory side, potential latency dependencies on frequency (Mäkelä et al., 2002; Roberts et al., 2000), and on the visual side, on eccentricity (Capilla et al., 2016; Eimer, 2000). Such lines of evidence demonstrate nonlinear dependencies on ERP latency, although shorter latencies are often associated with higher frequencies and, separately, with foveal locations.

For AH and AQ stimuli, AV precision conditions were separately controlled for the foreground and the background, so that these parameters did not match within any stimulus. This prevented participants from deploying a generic “Attend-All” strategy. Foreground and background AV precision conditions were counterbalanced across each attentional condition. To this end, the background precision of each stimulus was balanced according to a vector based on 5 background levels equally distributed across the AV precision range that never corresponded to the same level used for the foreground. For instance, if the foreground was set at ±99 ms asynchrony (level “2”, even), then the background could be either ±66, ±132, ±198, ±264, or ±330 ms (levels “1”, “3”, “5”, “7” or “9”, odd). Combination allocations were made by shuffling the level vector without replacement.

*Auditory stimulus probe*. This probe consisted of a 15-tone sequence created under the same specifications as the AV stimuli, presented without an accompanying visual stream. This auditory-only stimulus was used to detect reproducible auditory activity in the EEG (see *Data analysis*). The 15-tone sequence was quadruplicated and concatenated in time, lasting approximately 50 s.

**Task**

For each trial and participant, the combination of AV stimuli, the response keyword (‘more’ or ‘less’), and keyword left/right positions were randomized. To familiarize participants with each cue type and attentional condition, a practice phase consisting of 14 trials was introduced before the main session, with feedback given and with the experimenter present. Following the practice phase, participants were also asked to listen attentively and with their eyes closed to the auditory stimulus probe. During the main session, participants had an optional pause every 25 trials, and because responses were validated by a button press, participants had additional opportunities to pause. After the main session, the auditory stimulus probe procedure was repeated.

**Data analysis**

*EEG setup, preprocessing and denoising*. A 5^th^ order cascaded integrator-comb low-pass filter with -3 dB at 410 Hz was applied during recordings online, after which signals were decimated to 1024 samples per second. Online high-pass response was fully DC coupled. External recordings — including electrooculographic data — were taken supra- and infra-orbitally and from the left and right orbital rim plus the nasal tip. For offline denoising, single channel data were subsequently rejected in a blind manner according to a variance-based criterion (Junghöfer, Elbert, Tucker, & Rockstroh, 2000) that was applied for scalp sensor channels with a confidence coefficient $\lambda_{P}=4$. The procedure was repeated separately for the external reference channels. To reduce the effect of general movement artifacts, EEG data were submitted to independent component analyses using FastICA (Hyvarinen, 1999). Two independent components were automatically selected for their maximal proportion of broadband power in the 10-40 Hz region and projected out of the raw data. In a further procedure to reduce ocular artifacts, a time-shifted principal component analysis (de Cheveigné & Simon, 2007) was applied to discard environmental signals recorded on the oculogram reference sensors (shift: ±4 ms). A sensor noise suppression algorithm (de Cheveigné & Simon, 2008) was applied to attenuate artifact components specific to any single channel (63 neighbors). The blind variance-based rejection procedure was repeated resulting in less than 1% rejected single-channel trial time series on average (subject range 0.09% - 1.52%).

*DSS*. Pre- and post-experiment recordings from the “auditory stimulus probe” were equally pre-processed, but without downsampling, and epoched into 8 trials. To extract the reproducible DSS component, first, EEG data were bandpass-filtered with a second-order Butterworth filter in the 1 – 8 Hz region. To address variability across participants, which may affect spatial filter estimation, the blind rejection procedure described above was performed over channel and trial time series, including all participants’ data simultaneously and with a more restrictive confidence coefficient $\lambda_{P}=2.25$. The DSS component time series represents the most reproducible aspect of evoked responses and, similar to sensor selection in standard ERP studies, the component was selected as a single virtual sensor in all analyses.

*TRF estimation.* The linear model is formulated as:

$$r'\left( t \right)=\sum_{\tau} TRF\left( \tau\right)S\left( t-\tau\right)+\varepsilon\left( t \right)$$

where $\varepsilon\left( t \right)$ is the residual contribution to the evoked response not explained by the linear model and $TRF\left( \tau\right)$ represents the TRF under the conditions set by the chosen stimulus representation $S\left( t \right)$ for the same period of neural recording. This method therefore allowed us to investigate, for a given neural recording epoch $r\left( t \right)$, the auditory encoding of pips according to their association with the visuospatial task, thus we did not address the full auditory time series (including all pips) at once but selections thereof (e.g. $S_{VpA,{AH}_{ATT}}\left( t \right)$, see main text). For instance, the auditory stimulus time series of AQ trials were sub-divided to separately account for events associated to the target quadrant *AQ_ATT_* , versus its opposite quadrant *AQ_UAT_* , as well as the same abscissae and the same ordinates’ flanking quadrants *AQ_SAB_* and *AQ_SOR_*, respectively, e.g. $S_{ApV,AQ}\left( t \right)=S_{ApV,{AQ}_{ATT}}\left( t \right)+S_{ApV,{AQ}_{UAT}}\left( t \right)+S_{ApV,{AQ}_{SAB}}\left( t \right)+S_{ApV,{AQ}_{SOR}}\left( t \right)$ . For simplicity, in AQ conditions only target and opposite quadrants were considered in the main analysis.

For all three attentional conditions, single-trial data were organized according to AV precision as follows: a low AV precision group (‘Lo-AV’) that consisted of time-concatenated data from the bottom 5 AV precision levels; and a high AV precision group (‘Hi-AV’) corresponding to the top 5 levels. Concatenated EEG recordings were paired with corresponding concatenated stimuli time series.

*Statistical analysis*. To determine the presence of attentional effects, non-parametric tests where used where *t*-value time series were estimated from attended minus unattended TRF contrasts under a threshold corresponding to the distribution’s 0.05/8^th^ percentile, for randomization-based testing. After removal of Lo-AV conditions (see main text) the threshold was reset to the 0.05/4^th^ percentile. Any cluster exceeding the threshold was deemed significant if its associated *t*-statistic (sum of *t*-values within the cluster) exceeded those *t*-statistics estimated from the randomization distribution. The distribution was estimated after *N*=2^17^ resamplings, each generated by randomly shuffling between attended/unattended conditions per participant. To examine the interactions between precedence, spatial window, and temporal window, a three-way repeated measures ANOVA was employed.

**Supplementary Results**

*Behavioral results*. Participants’ global correct response rates in the 2-IFC task were significantly above chance level (mean ± SD: 67±9%; *t*(29)=4.3; *p*=1.8x10^-4^). Task performance did not significantly differ across the various foreground sizes (mean ± SD 70±14% Attend-All; 67±11% Attend-Half; 65±9% Attend-Quarter), as shown by a repeated-measures ANOVA of the participants’ performance scores for each attentional condition (*F*(2)=2.69; *p*=0.076).

*Attend-All TRFs*. In agreement with AH and AQ conditions, the resulting mean TRF across participants from AA trials reveals a biphasic response profile, with a positive and a negative peak in the 150-300 ms post-pip onset interval (Supplementary Figure 1B). This biphasic profile was preserved examined in relation to different temporal AV window precision and onset precedence order conditions (Supplementary Figure 1C). For Lo-AV conditions, the onset delay distributions have a median of ±128 ms (ApV and VpA respectively), interquartile range (IQR) ±128 ms, and range ±(0 - 330) ms. For Hi-AV, they have a median of ±39 ms, IQR ±56 ms, range ±(0 - 165) ms for Hi-AV (Figure 1C). Qualitatively, the TRF estimates show the early positive peak in the 150-200 ms interval, while a later negative peak in the 250-300 ms appears for tones closely followed by visual flips in the Hi-AV, ApV condition. Importantly, it is this condition where the distribution of flip presentations typically coincides with the sustained phase of tones.

*Control for Attend-Quarter stimuli duration*. Stimulus length differences between AH and AQ conditions could contribute to the observed cross-modal transfer results. To exclude the possibility that the significant effect observed for AQ trials but not AH may be driven by the relatively longer duration of the former (i.e. 30 pip-flip events in AH vs 60 in AQ) – and therefore by differential sustained attention drifting over time-, only data from the first half of each AQ trial were used for TRF estimation anew. Non-parametric permutation testing replicated the finding observed at ApV conditions, namely a significant difference by visuospatial attentional selection on the auditory TRF in the 262-324 ms post pip onset range (*p*=0.006). Repeated measures ANOVA on this duration-matched dataset revealed no effect of spatial window (F(1,27) = 0, p*=*1), an effect of precedence (F(1,27) = 6.123, p=0.020), with an precedence by spatial window interaction (F(1,27) = 4.654, p = 0.040). This last interaction was again due to half foregrounds showing no effect of precedence while quarter foregrounds do (F(1, 28) = 0.055, p = 0.817; and F(1,28) = 6.549; p = 0.016, respectively). The results confirm that the cross-modal attentional effects specific to quarter foregrounds were also sustained at these shorter periods.

*Control for Unattended Hi-AV TRFs.* AV precision differences between foreground and background sector presentations (see *Audiovisual stimuli*, above) could explain the observed cross-modal transfer results. To address this, additional TRFs were estimated from background pips of Hi-AV precision. These new TRFs were compared with the same Hi-AV foreground pip TRFs as before. Importantly, the procedure involved all Hi-AV presentations across the experiment (i.e. all epochs where both foreground *or* background are Hi-AV), thus extend beyond the subset of epochs where both foreground *and* background are Hi-AV. Non-parametric permutation testing corrected for multiple comparisons confirmed a significant effect of visuospatial attentional selection on the auditory TRF in the narrower 262-293 ms post pip onset range (*p*=0.033). Findings of greater peak amplitude for pips associated with ‘Attended’ than ‘Unattended Hi-AV’ visual sectors, were replicated on AQ, ApV trials (Figure 3D). A repeated-measures ANOVA of TRF contrasts in this window, with precedence (ApV vs. VpA) and spatial window (AH vs. AQ) as factors, showed no effect of spatial window (F(1,27) = 0, p*=*1), or precedence (F(1,27) = 2.386, p=0.134), and again confirmed a significant precedence by spatial window interaction (F(1,27) = 5.323, p = 0.029). As before, this interaction was due to the half foregrounds showing no effect of precedence, while this effect was significant for the quarter foregrounds (F(1, 28) = 1.411, p = 0.245; and F(1,28) = 6.639; p = 0.016, respectively). Given that Hi-AV background TRFs were obtained regardless of correspondence to the same epochs as Hi-AV foreground TRFs, these findings suggest that changes to auditory representations by visuospatial association may occur independently of the particular foreground-background dynamics on a given trial.

*Control for alternative AV matches*. Alternative pip-flip associations could be formed in the event that a Lo-AV background pip happens to occur more closely in time than with a foreground flip. To address the involvement of systematic associations that differ from the nominal pairings, AQ TRFs were re-estimated with pip-flip pairings based on temporal rank order, regardless of frequency and eccentricity association with flips. Non-parametric permutation testing corrected for multiple comparisons confirmed the significant effect of visuospatial attentional selection previously observed in the 262-293 ms post pip onset range (*p*=0.015). A repeated-measures ANOVA of TRF contrasts in this window, with precedence (ApV vs. VpA) as factor revealed a significant effect (F(1,26) = 9.414, p=0.005) confirming that cross-modal transfer was instantiated by pip onset precedence.

**Supplementary Figures**

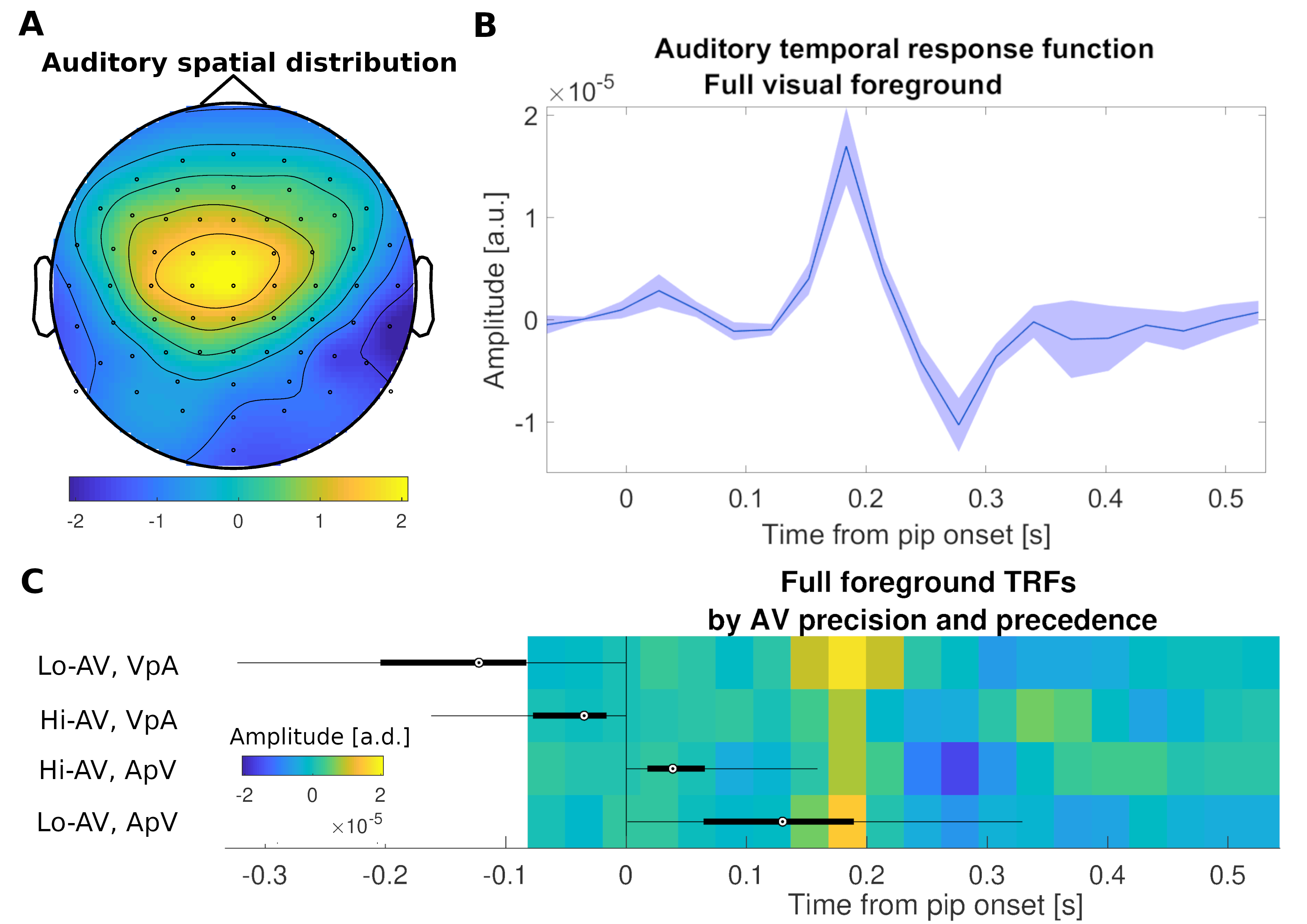

**Supplementary Figure 1.** *Auditory encoding of pip sounds during concurrent visuospatial selection*. (A) EEG topography of top reproducible auditory activity across participants. Recordings were projected onto this component via spatial filtering for TRF estimation. (B) TRF grand-average from all Attend-All trials (full foreground). The auditory encoding of pips in AV presentations results in a biphasic profile in the TRF. Surface area corresponds to ±1 standard error of the mean. (C) TRFs according to AV precision and precedence order. Four grand average TRFs, with amplitudes now depicted by depth, represent encoding specific to Hi- versus Lo-AV precision conditions. Individual pips are also separated by precedence order (ApV or VpA). Boxplots indicate the temporal distribution corresponding to flip onset times. The earlier positive TRF peak, which appears locked to the pip onset, can be observed across all conditions. The late negative TRF peak appears most clearly in Hi-AV ApV condition, as flips occur early on during ongoing pip processing along with the sustained phase of the sound.

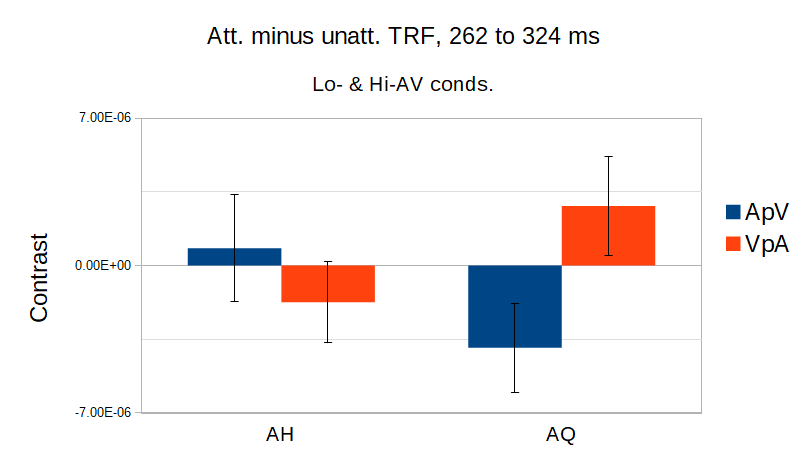

**Supplementary Figure 2.** *Significant auditory precedence by visual foreground size interaction*. Results presented in a form relevant for interpreting the interaction of precedence (ApV vs. VpA) and foreground size (AH vs. AQ). ApV order induces greatest attentional contrast change at quarter but not half foreground sizes. Error bars represent 1 SEM.

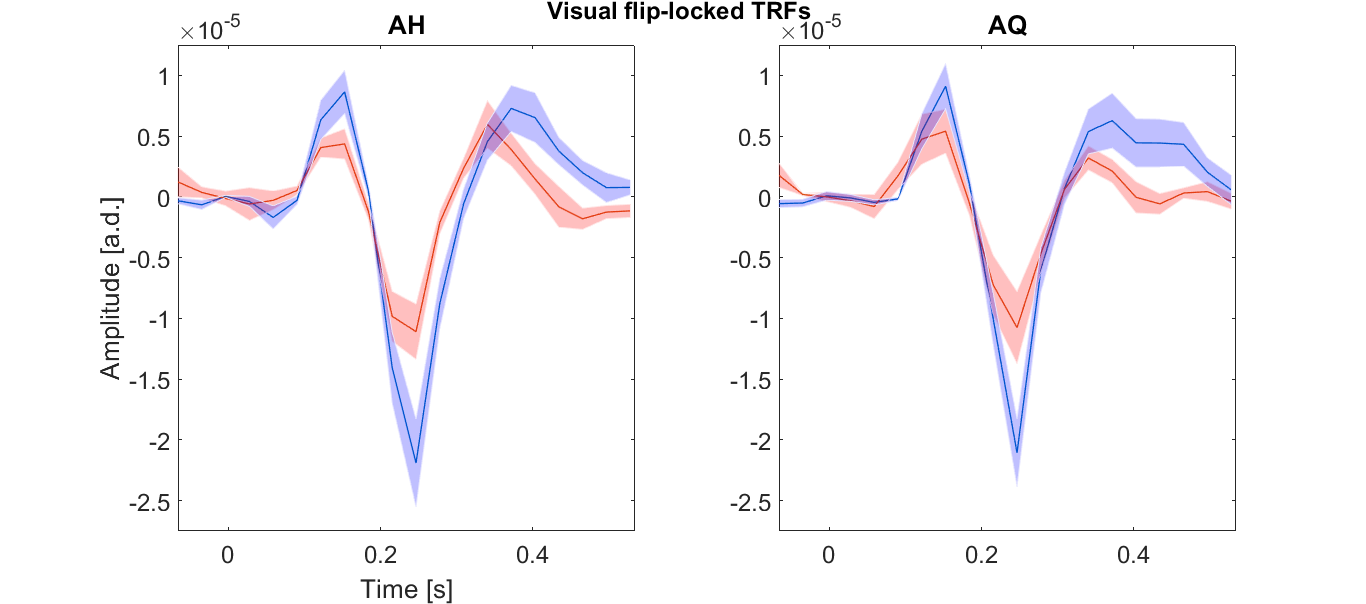
**Supplementary Figure 3.** *Visual flip-locked temporal response functions*. Flips from attended (blue) versus unattended visual sectors (red) exhibit differential encoding by selection across conditions. AH: significant clusters found in the 246-309 ms and 403-496 ms intervals (*p*=2.5x10^-4^ and *p*=5.6x10^-4^ respectively). AQ: significant clusters found in the 246-278 ms and 403-465 ms intervals (*p*=0.011 and *p*=0.006, respectively).

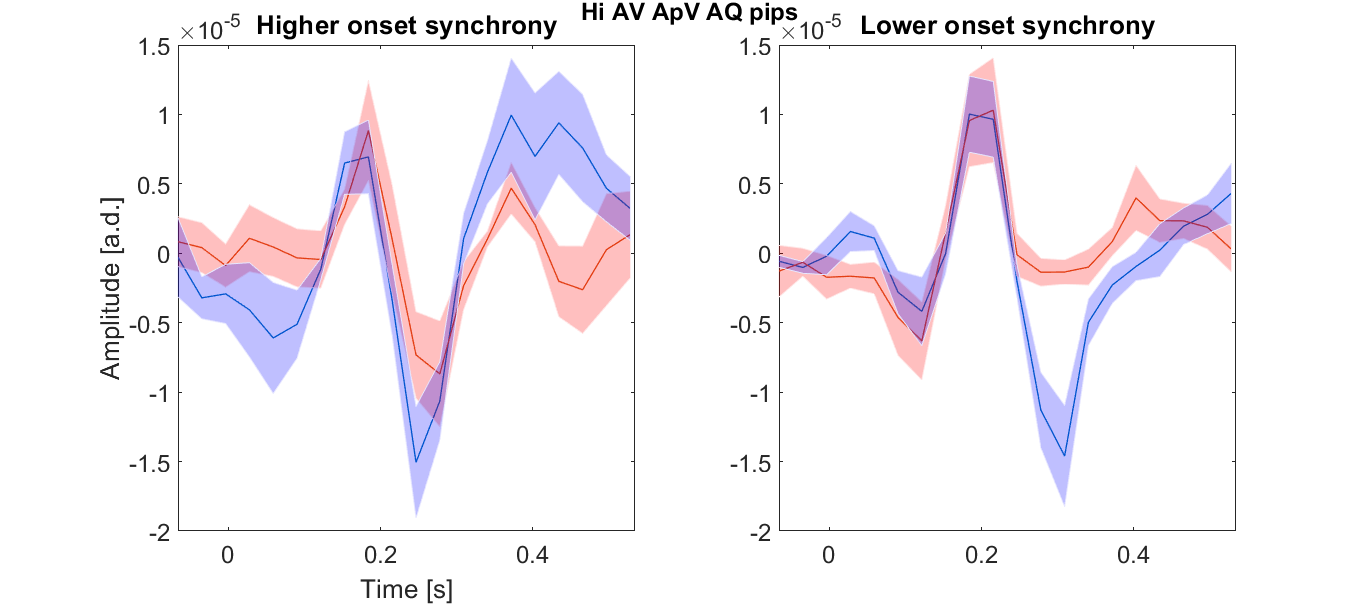
**Supplementary Figure 4.** *Pip temporal response functions from higher and lower synchrony pips*. These pips from AQ, ApV conditions were separated by a median split procedure on their onset asynchrony with respect to corresponding flips. Pips with higher synchrony within this set (<39 ms) did not show a significant attentional effect cluster (left). By contrast, for pips within the lower synchrony range (39 – 165 ms range) did replicate the cross-modal transfer effect, in the form of a significant cluster between pips associated to the attended (blue) and unattended (red) visual sector in the 262-324 ms window (*p*=7.5 x 10^-4^, right).

***Supplementary References***

Capilla, A., Melcón, M., Kessel, D., Calderón, R., Pazo-Álvarez, P., & Carretié, L. (2016). Retinotopic mapping of visual event-related potentials. *Biological Psychology*, *118*, 114–125. doi: 10.1016/j.biopsycho.2016.05.009

de Cheveigné, A., & Simon, J. Z. (2007). Denoising based on time-shift PCA. *Journal of Neuroscience Methods*, *165*(2), 297–305. doi: 10.1016/j.jneumeth.2007.06.003

de Cheveigné, A., & Simon, J. Z. (2008). Sensor noise suppression. *Journal of Neuroscience Methods*, *168*(1), 195–202. doi: 10.1016/j.jneumeth.2007.09.012

Eimer, M. (2000). An ERP study of sustained spatial attention to stimulus eccentricity. *Biological Psychology*, *52*(3), 205–220. doi: 10.1016/S0301-0511(00)00028-4

Hyvarinen, A. (1999). Fast and robust fixed-point algorithms for independent component analysis. *IEEE Transactions on Neural Networks*, *10*(3), 626–634. doi: 10.1109/72.761722

Junghöfer, M., Elbert, T., Tucker, D. M., & Rockstroh, B. (2000). Statistical control of artifacts in dense array EEG/MEG studies. *Psychophysiology*, *37*(04), 523–532. doi: null

Mäkelä, A. M., Alku, P., Mäkinen, V., Valtonen, J., May, P., & Tiitinen, H. (2002). Human cortical dynamics determined by speech fundamental frequency. *NeuroImage*, *17*(3), 1300–1305.

Roberts, T. P., Ferrari, P., Stufflebeam, S. M., & Poeppel, D. (2000). Latency of the auditory evoked neuromagnetic field components: Stimulus dependence and insights toward perception. *Journal of Clinical Neurophysiology: Official Publication of the American Electroencephalographic Society*, *17*(2), 114–129. doi: 10.1097/00004691-200003000-00002
